## Supplementary Figures; legends for Supplementary Tables for "Ribosome profiling of porcine reproductive and respiratory syndrome virus reveals novel features of viral gene expression"

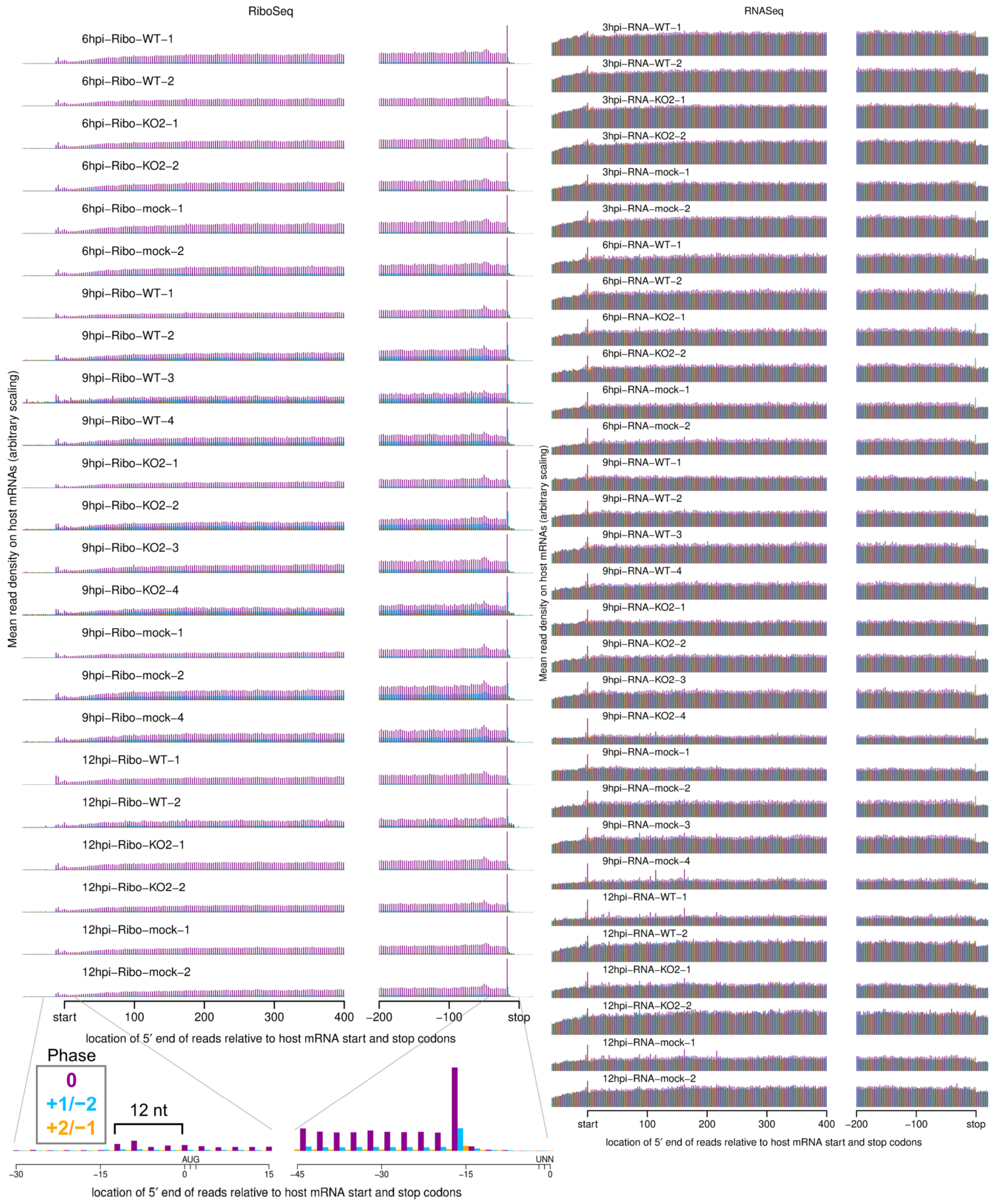


**Supplementary Figure 1. Metagene profile showing the average distribution of 5′ ends of host mRNA-mapping reads relative to start and stop codons.**

Only transcripts with an annotated CDS of at least 150 codons, 5′ UTR of at least 60 nt, and 3′ UTR of at least 90 nt were included in the analysis. The total number of positive-sense reads from all these transcripts mapping to each position was plotted. RiboSeq reads originate from translating ribosomes and therefore display clear triplet periodicity (phasing), with few reads mapping to the UTRs, and a heightened termination peak characteristic of samples harvested without CHX pre-treatment. Underneath the 12hpi-Ribo-mock-2 library, a magnified view of 45 nucleotides around the start and stop codons for that library is shown, with the typical 12-nt distance between the 5′ end of RPFs and the ribosomal P site indicated for initiating ribosomes. This +12 nt offset is applied to read 5′ end coordinates to plot reads at inferred ribosomal P site positions in all plots except this and similar plots in Supplementary Figure 6B and Supplementary Figure 12D. RNASeq reads do not originate from ribosomal protection, and therefore display a roughly uniform distribution with no clear dominant phase.

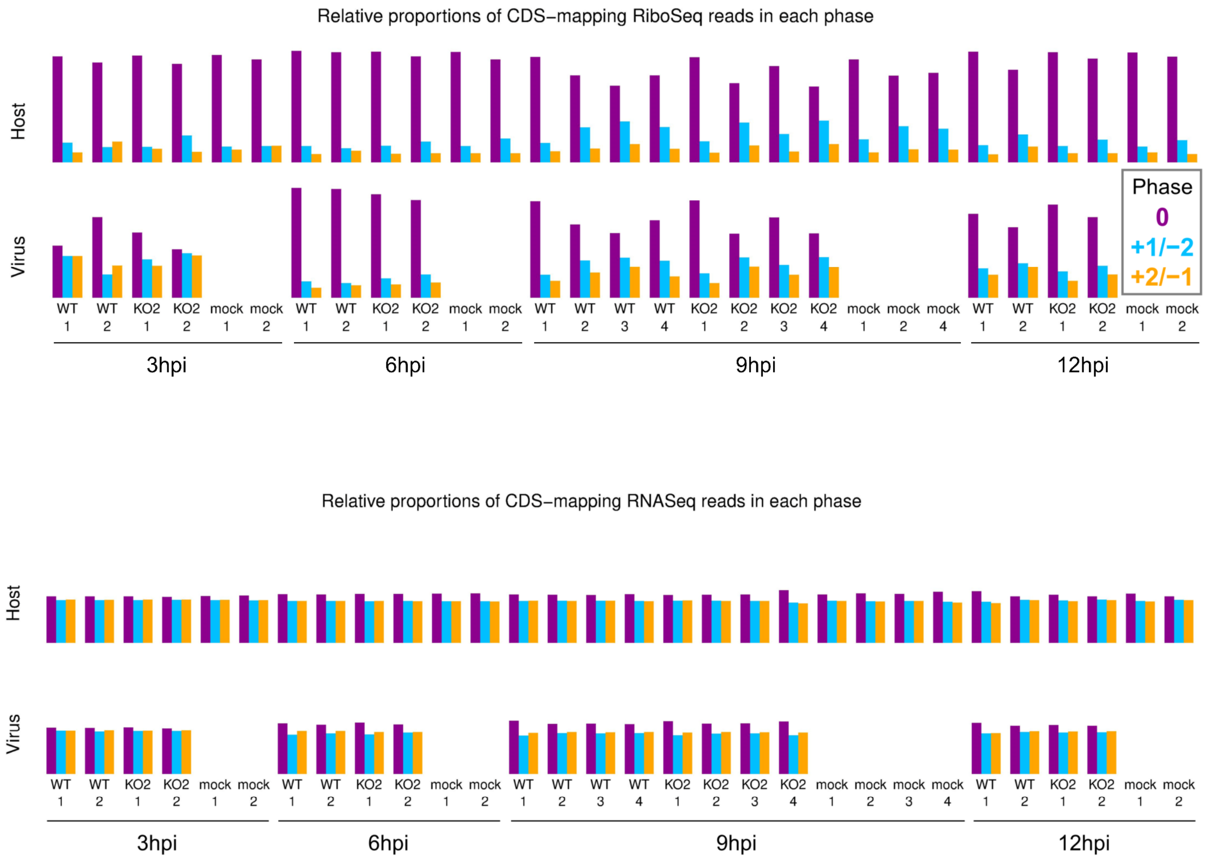


**Supplementary Figure 2. Phase composition of CDS-mapping reads.**

**Upper:** Proportion of RiboSeq reads (all read lengths) attributed to each phase, from positive-sense reads mapping within host (top) or viral (bottom, mock excluded) CDSs. Overlapping regions of viral CDS were excluded. 3 hpi libraries are shown here to provide an additional method to verify that the virus-mapping reads have a relatively low proportion of genuine RPFs, hence exclusion of these libraries from all other quality control analyses. **Lower:** Analysis from the upper panel carried out on RNASeq reads.

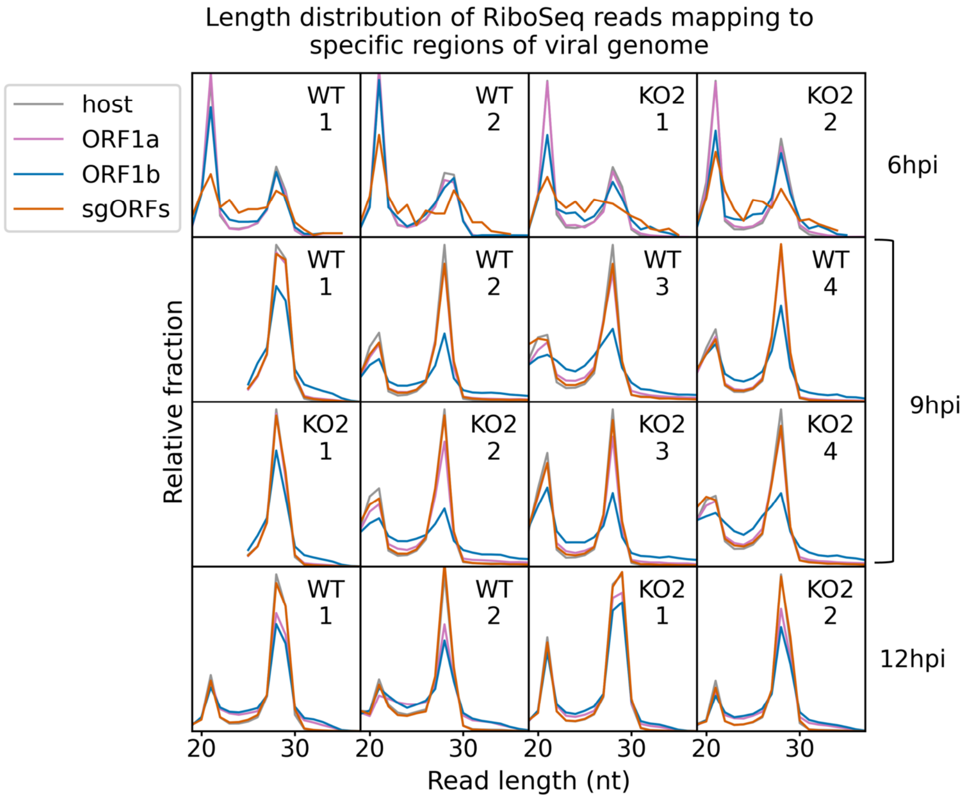


**Supplementary Figure 3. Length distribution of RPFs mapping to specified regions of the viral genome.**

Only positive-sense RiboSeq reads were used. Reads mapping to ORF1a are shown in purple, ORF1b in blue, and sgRNA ORFs in orange. The length distribution of host CDS-mapping RPFs from Figure 1C is reproduced in grey for comparison. Note that read counts for each length were normalised by the total number of reads in the region to make the scales comparable between regions. The coordinates used to define each region of the viral genome can be found in Supplementary Table 1. Note that, as sgRNA transcription and translation is minimal at 6 hpi, very few (156–293) reads formed the input for the sgORF-mapping read length distributions at this timepoint, and these are therefore likely highly subject to noise.

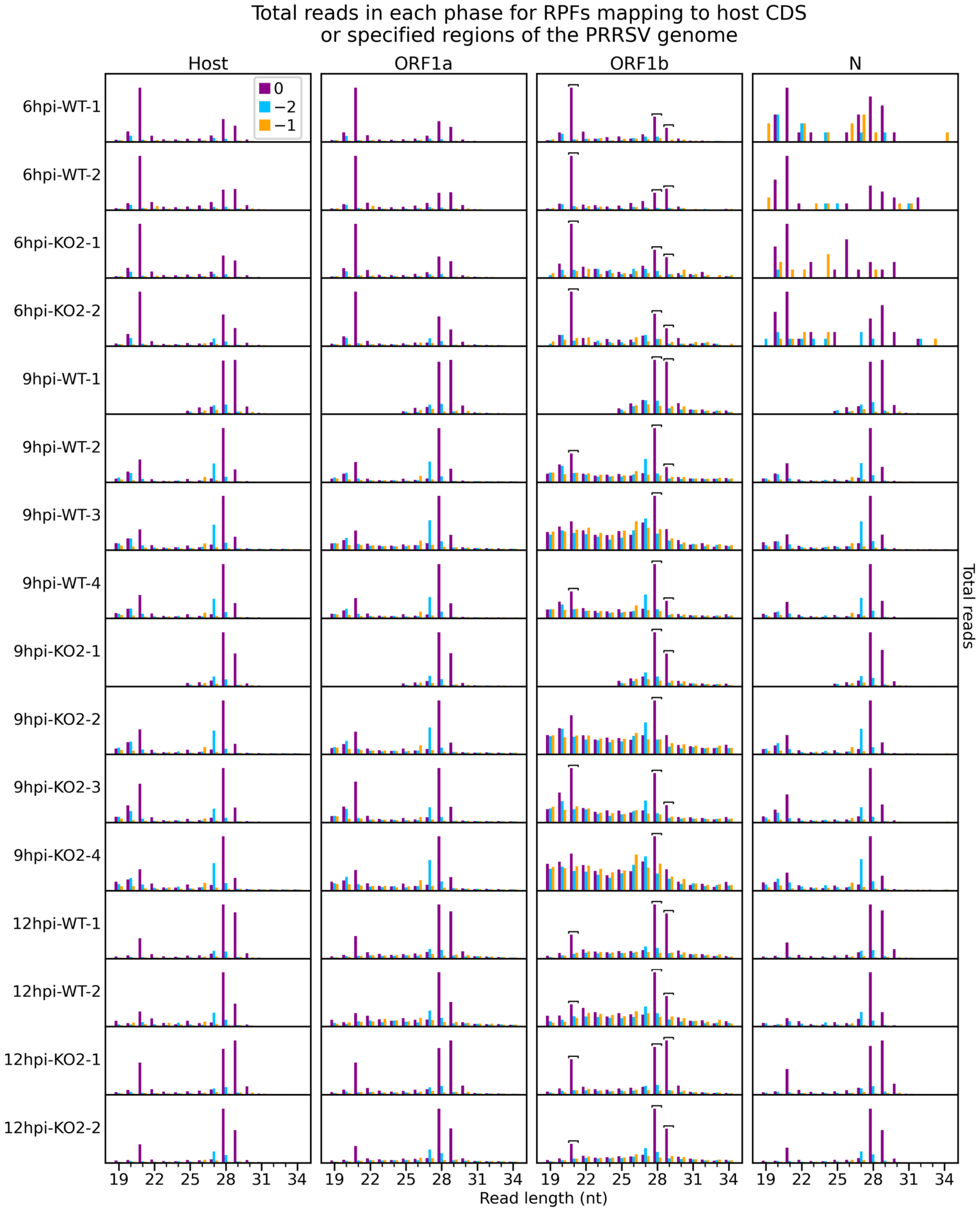


**Supplementary Figure 4. Number of reads of each length that are attributed to each phase, for RPFs mapping to specified regions of the viral genome.**

Only positive-sense RiboSeq reads were used, and regions from Supplementary Figure 3 were edited to exclude overlapping CDSs (coordinates given in Supplementary Table 1). Phase 0 is designated independently for each region, relative to the first nucleotide of that region's ORF, to aid comparison (e.g. the dominant phase in ORF1b is designated 0 instead of −1). The host phasing information from Supplementary Figure 2 was stratified by read length and reproduced for comparison. Note that y-axis scaling was set for each library and region separately to facilitate comparison. Read lengths forming the peaks of the characteristic bimodal RPF length distribution in this dataset (21, 28 and 29 nt) were selected as likely having the highest signal to noise ratio. From within this selection, read lengths which show similar phasing in the ORF1b region of the viral genome and host CDSs were determined for each library individually and designated as showing minimal RNP contamination. These are indicated by square brackets, and are used for all analyses involving these libraries unless specified. As for Supplementary Figure 3, very few (34–49) reads formed the input for the 6 hpi N phase composition plots, and these are therefore likely highly subject to noise.

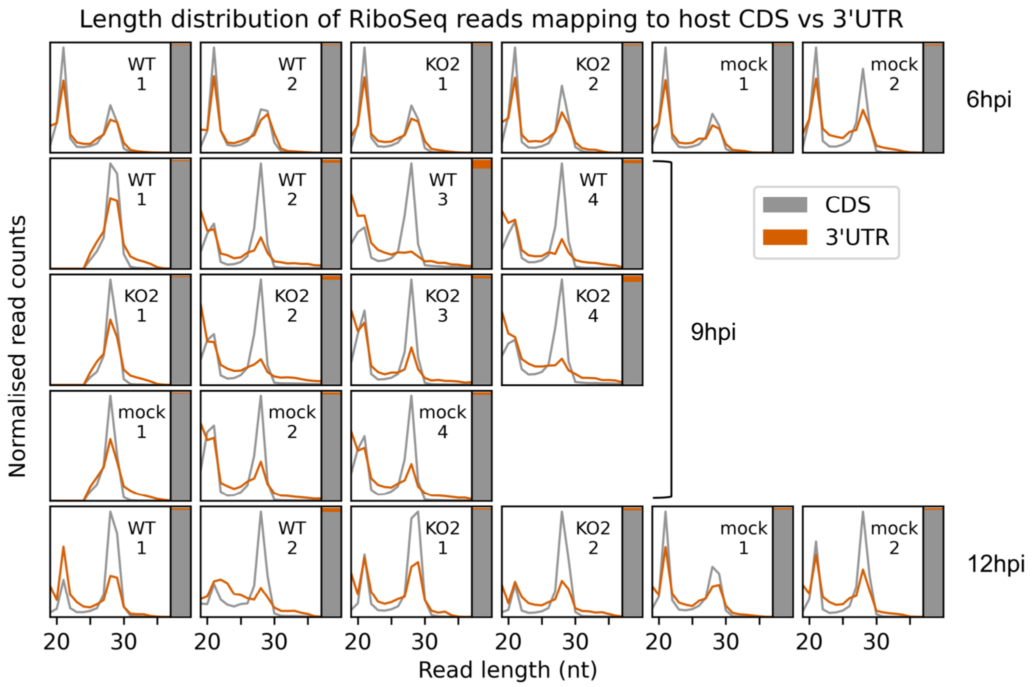


**Supplementary Figure 5. Assessment of potential RNP contamination of host-mapping RiboSeq reads.**

Length distribution (line graph) and relative density (stacked bar chart) of RiboSeq reads mapping to host CDSs (grey) compared to host 3′ UTRs (orange). Only host mRNAs with a CDS of at least 150 codons and a 3′ UTR of at least 100 codons were included in the analysis. The length distributions of positive-sense RPFs with inferred P sites in the CDS (codons −100 to −10 relative to the stop codon) and 3′ UTR (codons +10 to +100 relative to the stop codon) were determined and normalised by the total number of reads in each category to make the scales comparable. To the right of each length distribution, a stacked bar chart indicates the relative density of reads in each region (note that the 3′ UTR in orange is a minor component in this chart, reflective of its low read density relative to the CDS). RNP contamination is expected to affect both the CDSs and the UTRs but would be more visible in the UTRs due to the lower density of genuine RPFs (largely originating from multi-mapping reads aligning to transcript variants with identical regions annotated as CDS for some isoforms and 3′ UTR for others; multi-mapping reads were randomly assigned to a unique site). The uncontaminated 3′ UTR read length distribution should match the CDS-mapping read length distribution, but with lower relative density, as observed here in most samples. Where there are differences they are also present in mock libraries, indicating they are not related to infection. In any case, the relative density of 3′ UTR-mapping reads remains very low, indicating RNP contamination is unlikely to affect most host mRNA analyses.

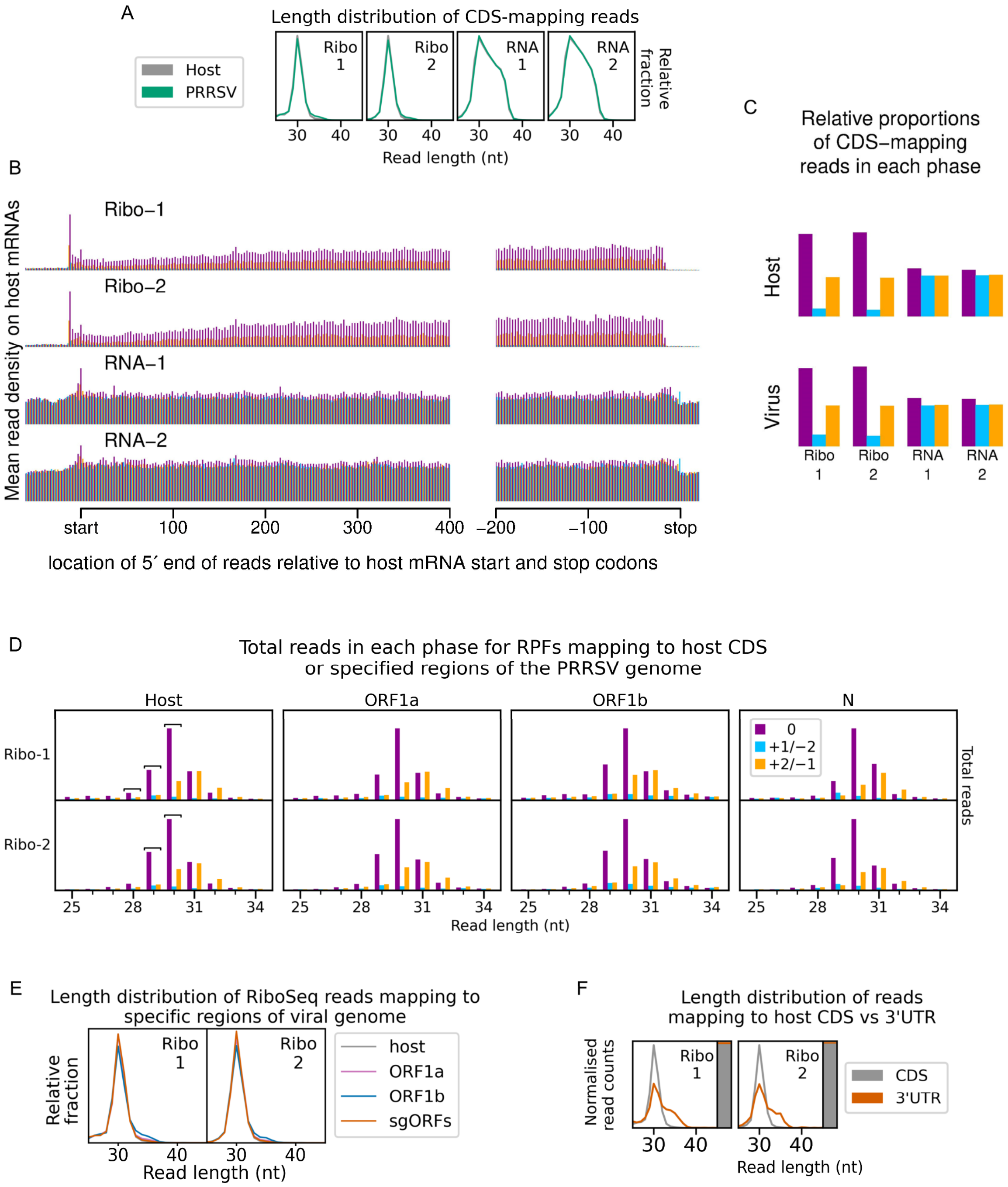


**Supplementary Figure 6. Quality control for the EU PRRSV dataset.**

**A)** Length distribution of positive-sense host (grey) and virus (green) reads mapping within CDSs. Fragments of 25–34 nt were size-selected during the library preparation for all samples. **B)** Metagene profile of the average distribution of 5′ ends of host mRNA-mapping reads relative to start and stop codons. Plot constructed as in Supplementary Figure 1. **C)** Proportion of host and viral reads attributed to each phase. Plot constructed as in Supplementary Figure 2. **D)** Number of reads of each length that are attributed to each phase, for reads with 5′ ends mapping to specified regions of the viral genome. Plot constructed as in Supplementary Figure 3. RNP contamination in these libraries is minimal, but to aid visualisation of phasing, read lengths for which at least two thirds of host CDS-mapping reads are in phase 0 were selected for some (specified) plots and analyses. These read lengths are indicated by square brackets. **E)** Length distribution of positive-sense RiboSeq reads with 5′ ends mapping to specified regions of the viral genome: ORF1a (purple), ORF1b (blue), sgRNA ORFs (orange). The length distribution of host CDS-mapping RPFs from A is reproduced for comparison (grey). The coordinates used to define each region of the viral genome for this and panel D can be found in Supplementary Table 1. **F)** Length distribution (line graph) and relative density (stacked bar chart) of reads mapping to host CDSs (grey) compared to host 3′ UTRs (orange). Plot constructed as in Supplementary Figure 5.

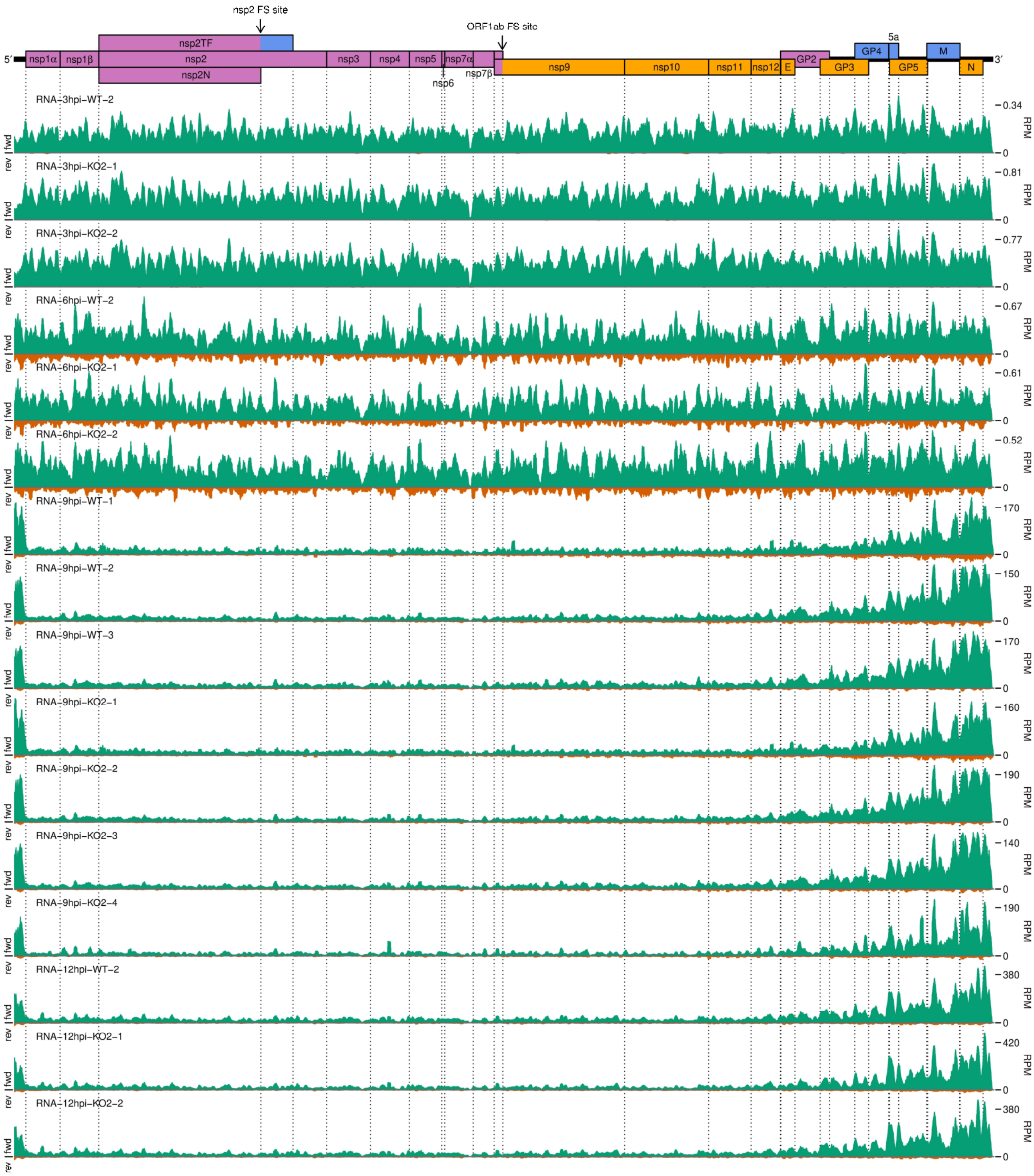


**Supplementary Figure 7. Further replicates of a timecourse of NA PRRSV viral transcription.**

RNASeq read densities on the viral genome, for all libraries not shown in Figure 2B. Plots constructed as in Figure 2B.

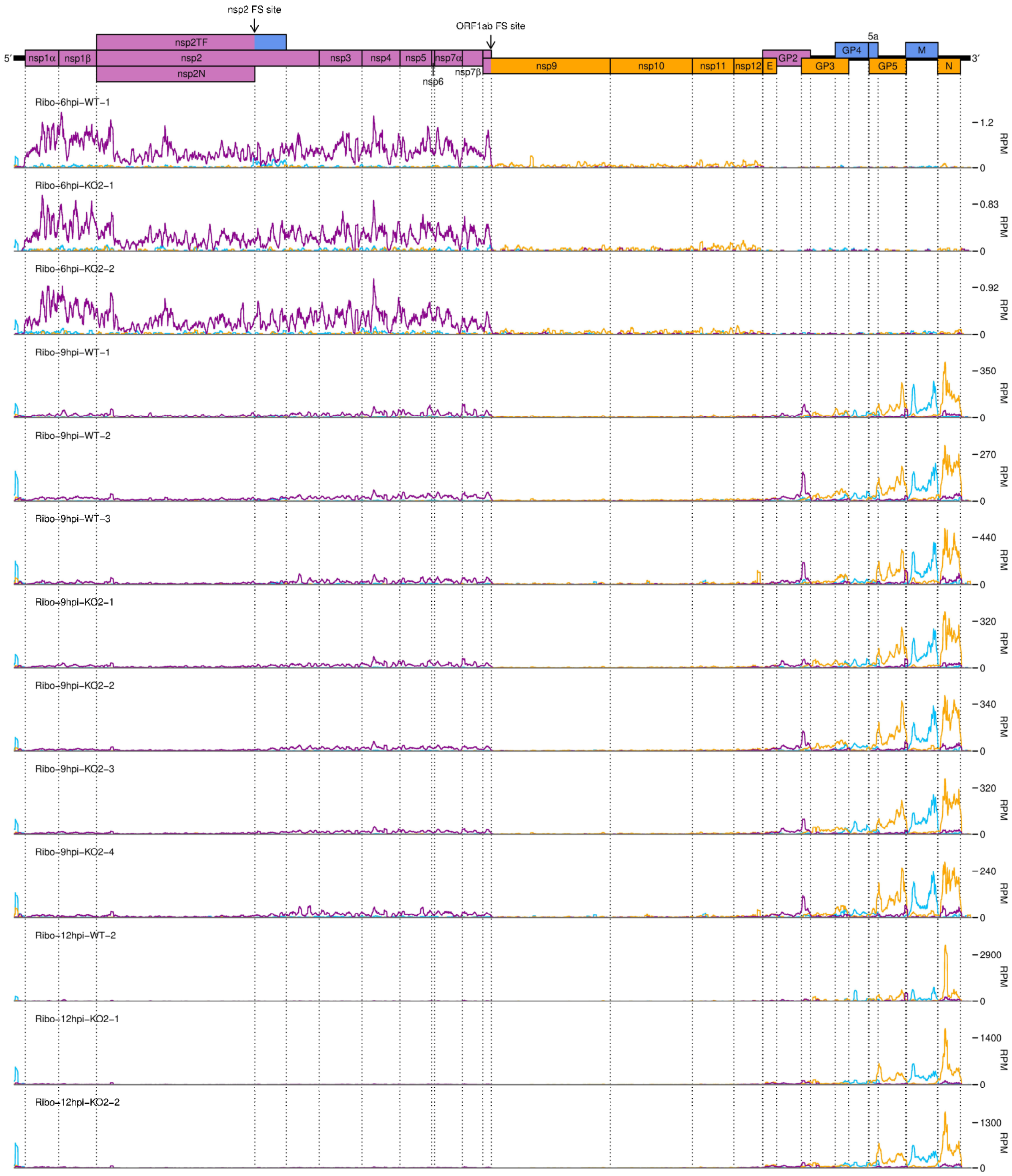


**Supplementary Figure 8. Further replicates of a timecourse of NA PRRSV viral translation.**

RiboSeq read densities on the viral genome, for all libraries not shown in Figure 2C. Plots constructed as in Figure 2C.

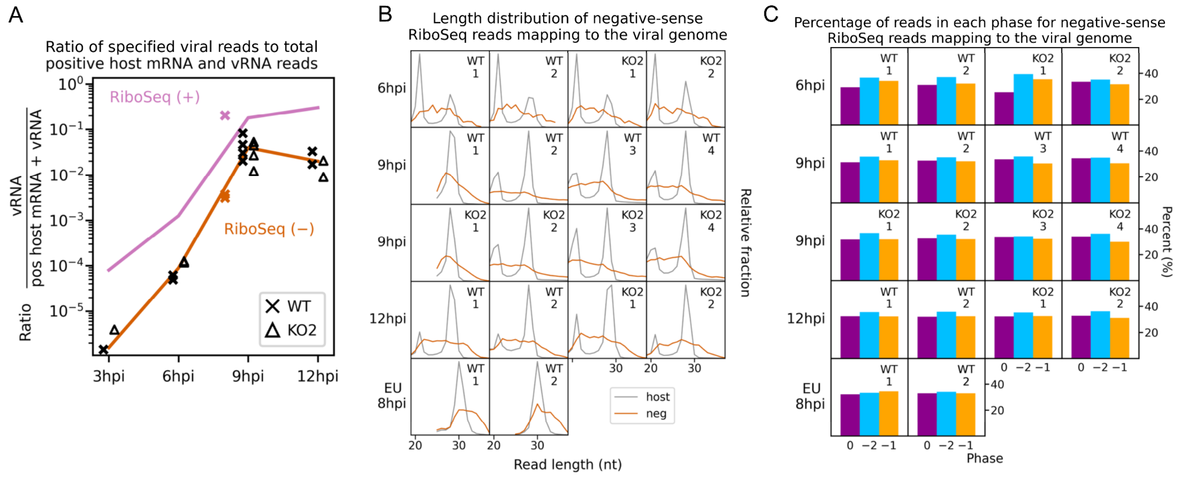


**Supplementary Figure 9. Investigation of negative-sense RiboSeq reads.**

**A)** Ratio of virus-mapping reads to [positive-sense host mRNA- plus positive-sense vRNA-mapping reads] for negative-sense RiboSeq reads. Mean values for RiboSeq (+) at each timepoint for NA PRRSV (lines) and individual datapoints for EU PRRSV (crosses) are reproduced from Figure 2D for comparison. Plot constructed as in Figure 2D. **B)** Length distribution of negative-sense RiboSeq reads (orange) mapping to the viral genome for NA (6 to 12 hpi) or EU (8 hpi) PRRSV. The length distribution of host CDS-mapping RPFs from Figure 1C and Supplementary Figure 6A is reproduced for comparison (grey). **C)** Percentage of reads attributed to each phase for negative-sense RiboSeq reads mapping to the viral genome for NA (6 to 12 hpi) or EU (8 hpi) PRRSV.

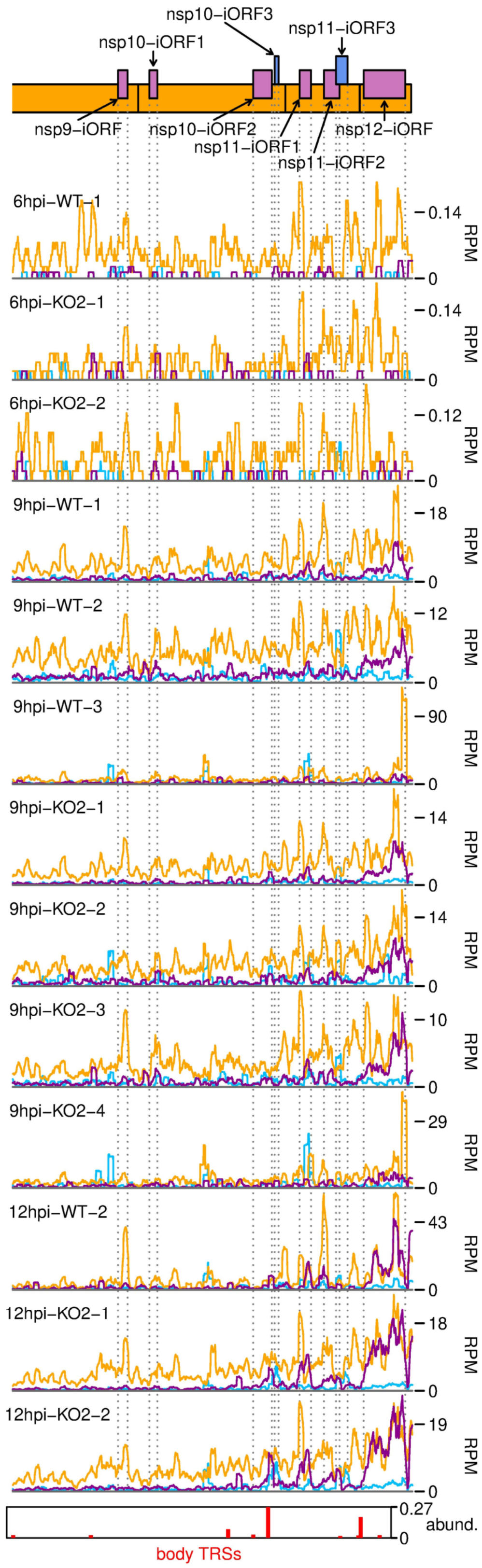


**Supplementary Figure 10. Further replicates of RPFs mapping to novel ORFs overlapping ORF1b.**

Plot constructed as in Figure 6B.

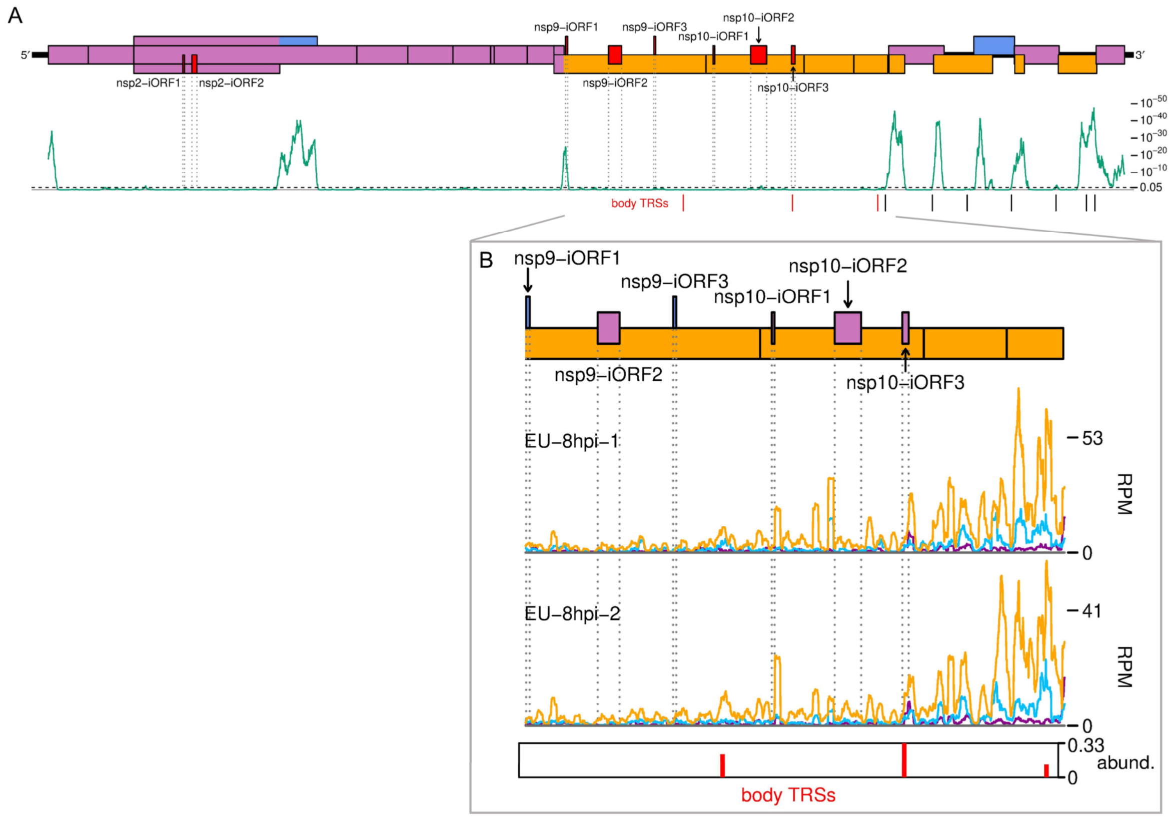


**Supplementary Figure 11. The EU PRRSV translatome.**

**A)** Locations of novel ORFs in the EU PRRSV genome. Plot constructed as in Figure 6A, with synonymous site conservation analysed for 120 available EU PRRSV sequences, and positions of body TRSs within ORF1b shown in red. **B)** Translation of EU PRRSV novel ORFs overlapping ORF1b. ORF1b is shown in its entirety, and the plot is constructed as in Figure 6B, with body TRSs with over 10 junction-spanning reads included, and body TRS bar heights representing junction-spanning reads (for both libraries combined) scaled relative to the canonical sgRNA with the fewest junction-spanning reads (GP3). Only read lengths with good phasing (indicated in Supplementary Figure 6D) were used to generate this plot.

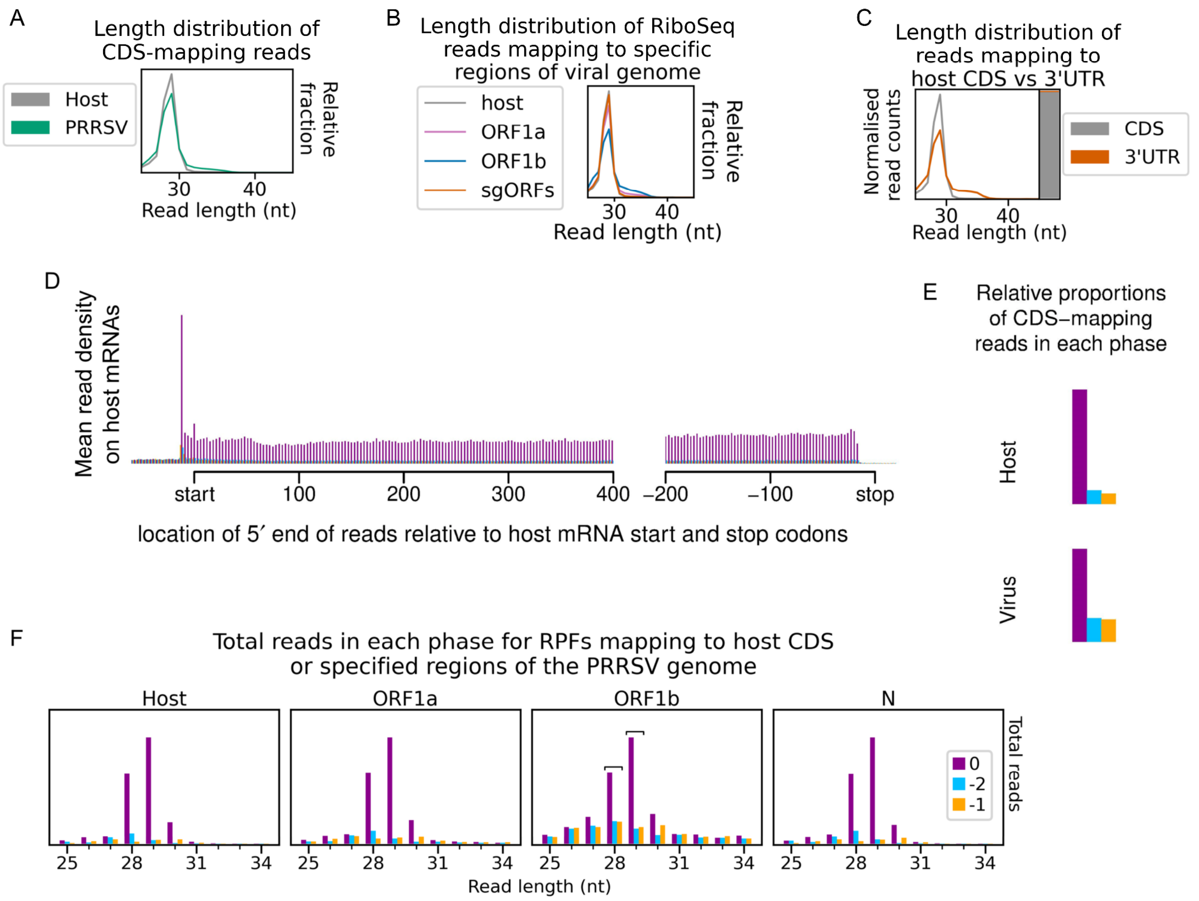


**Supplementary Figure 12. Quality control for NA PRRSV CHX dataset.**

**A)** Length distribution of positive-sense host (grey) and virus (green) reads mapping within CDSs. Fragments of 25–34 nt were size-selected during the library preparation. **B)** Length distribution of positive-sense RiboSeq reads with 5′ ends mapping to specified regions of the viral genome: ORF1a (purple), ORF1b (blue), sgRNA ORFs (orange). The length distribution of host CDS-mapping RPFs from A is reproduced for comparison (grey). The coordinates used to define each region of the viral genome for this and panel F can be found in Supplementary Table 1. **C)** Length distribution (line graph) and relative density (stacked bar chart) of reads mapping to host CDSs (grey) compared to host 3′ UTRs (orange). Plot constructed as in Supplementary Figure 5. **D)** Average distribution of 5′ ends of host mRNA-mapping reads relative to start and stop codons. Plot constructed as in Supplementary Figure 1. **E)** Proportion of host and viral reads attributed to each phase. Plot constructed as in Supplementary Figure 2. **F)** Number of reads of each length that are attributed to each phase, for reads with 5′ ends mapping to specified regions of the viral genome. Plot constructed as in Supplementary Figure 4. Read lengths selected as showing minimal effect of RNP contamination are indicated by square brackets, and are used in all plots and analyses for this library.

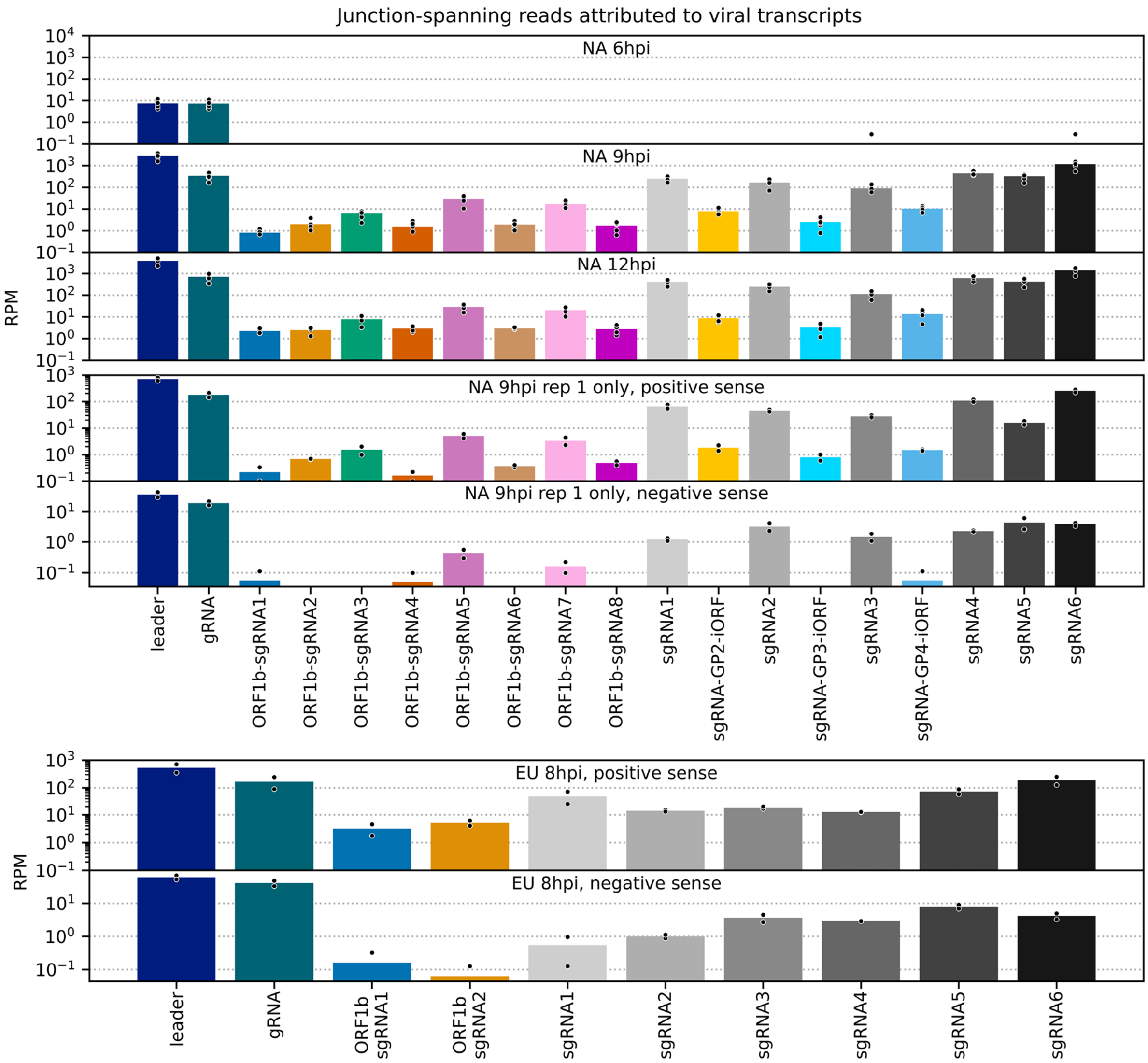


**Supplementary Figure 13. Number of junction-spanning reads attributed to viral transcripts.**

Data for canonical transcripts are reproduced from Figure 8A, and the same junction-spanning read abundance calculations were performed for novel transcripts. Bars represent the mean, with individual data points represented as black circles with white outlines. Datapoints with values of zero were excluded from the scatter plot but included in the mean calculations.

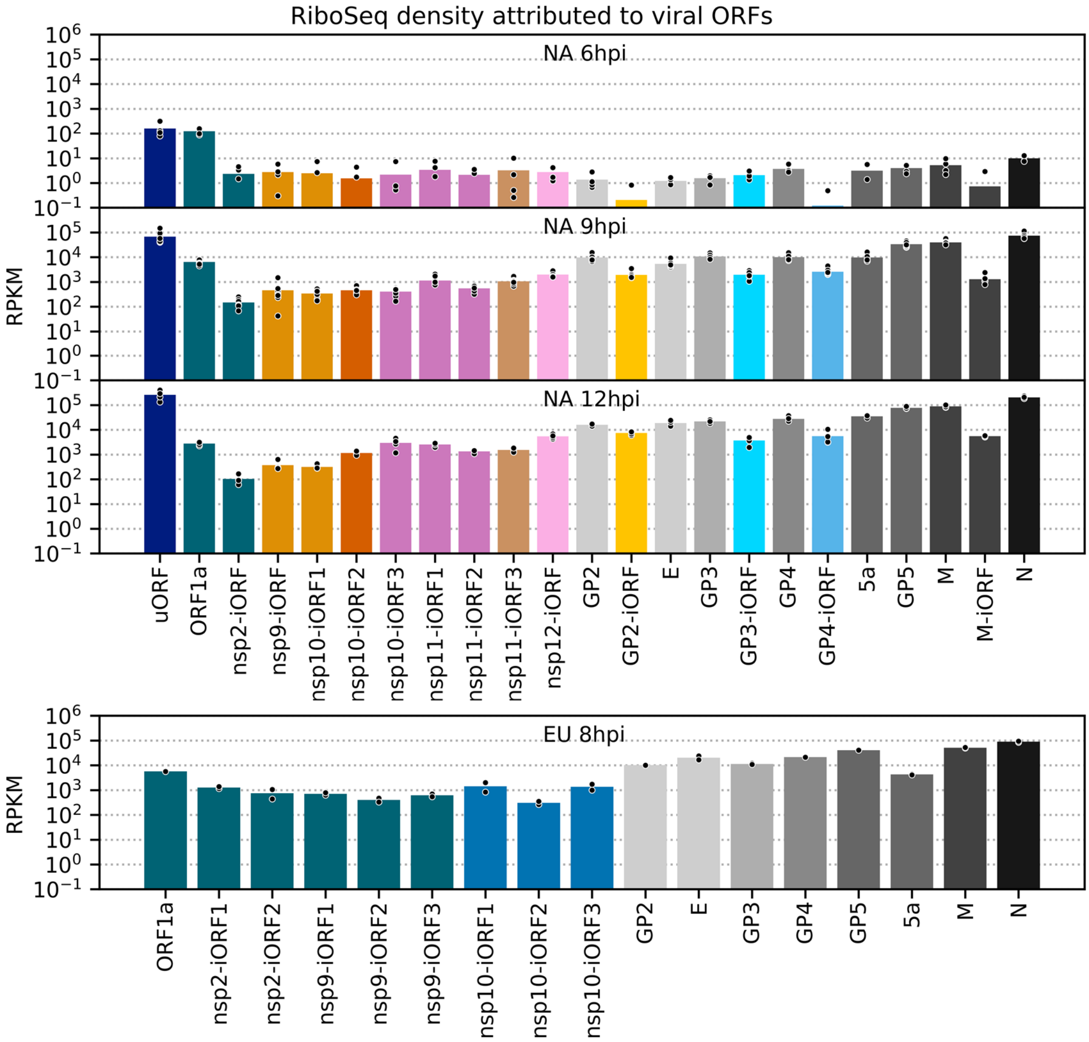


**Supplementary Figure 14. RiboSeq density attributed to each viral ORF.**

Data for canonical ORFs is reproduced from Figure 8A, and the same calculations were performed for novel ORFs. Plot constructed as in Supplementary Figure 13, with ORFs coloured according to their paired transcript (pairings in Figure 7, colour legend in Figure 8).

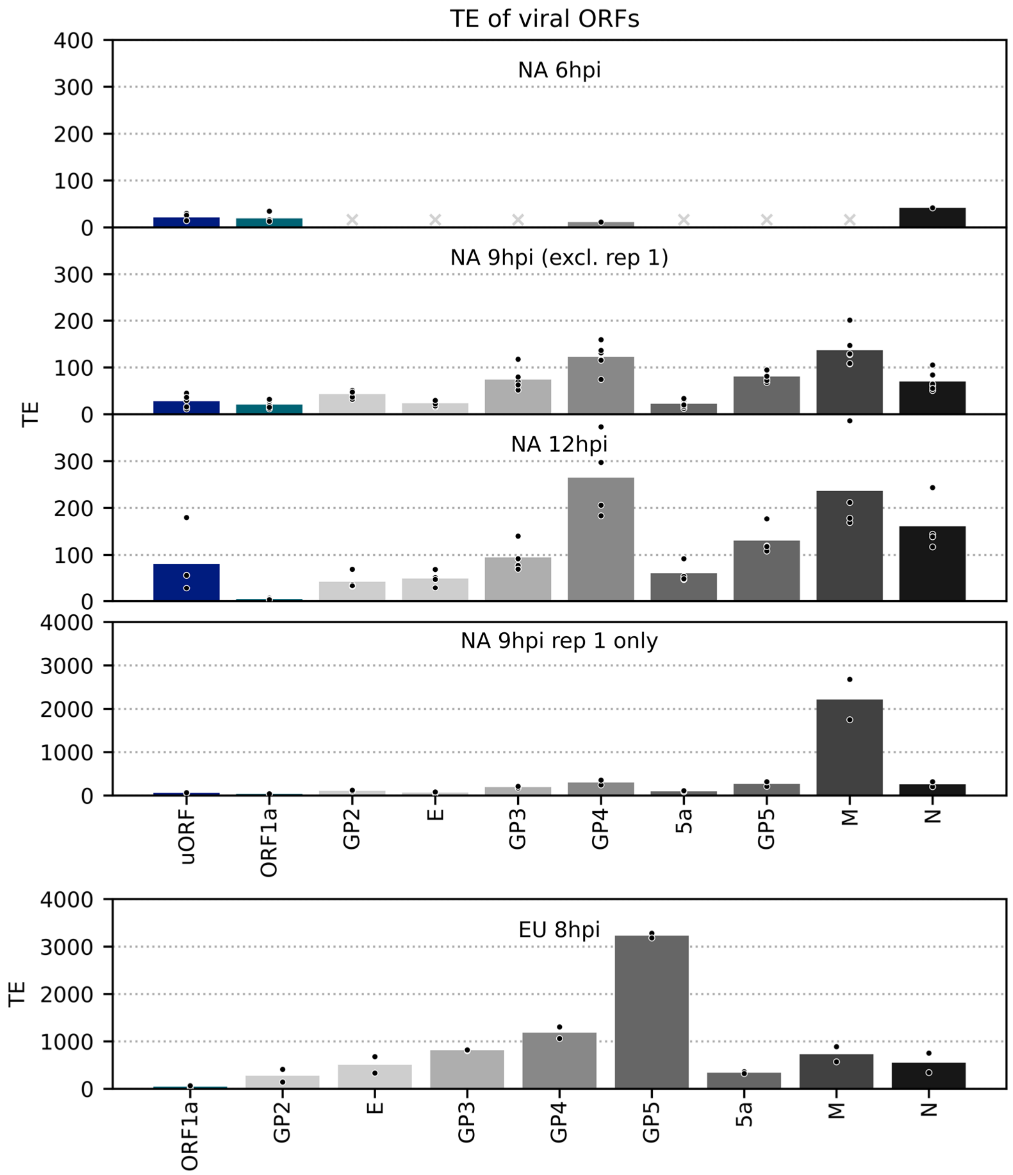


**Supplementary Figure 15. Translation efficiency of viral ORFs.**

TE, calculated as (RPF density in RPKM) / (junction-spanning reads in RPM), of canonical viral ORFs and the NA PRRSV uORF. Other novel ORFs were excluded, as this method can be unreliable for small, lowly expressed overlapping ORFs and/or lowly abundant transcripts due to greater susceptibility to noise. In cases where no junction-spanning reads were detected, TE was not calculated, with grey crosses in place of bars indicating that this was the case for all libraries within a group. Plot constructed as in Supplementary Figure 13, using a linear scale and with ORFs coloured according to their paired transcript. The high estimated TE of M for the 9 hpi replicate one libraries likely reflects the sensitivity of sgRNA 5 transcript abundance estimation to the shorter read lengths in these libraries (Figure 8B, Supplementary Figure 13).

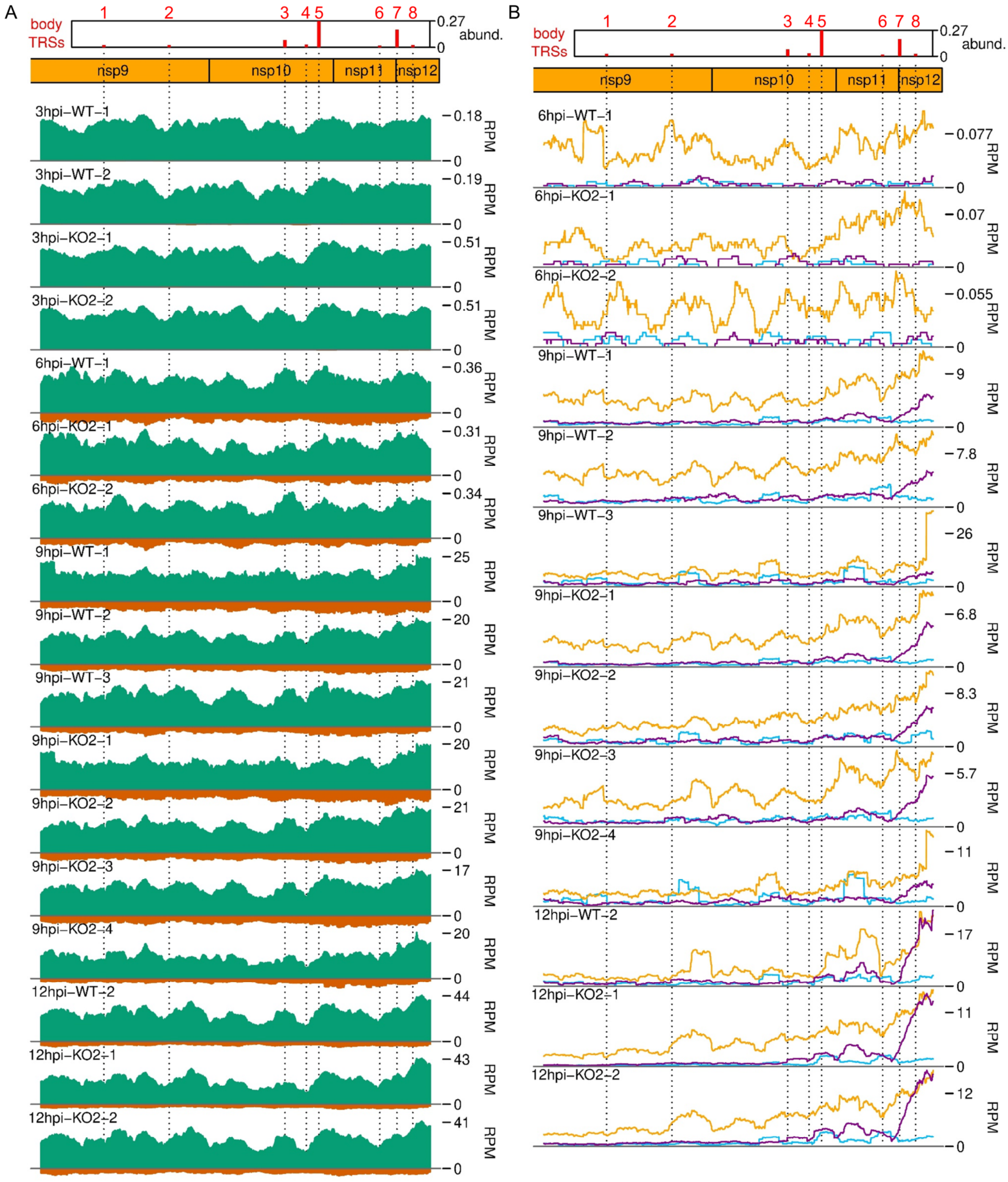


**Supplementary Figure 16. Distribution of A) RNASeq and B) RiboSeq reads mapping to the ORF1b region of the NA PRRSV genome, further replicates.**

Plot constructed as in Figure 9A.

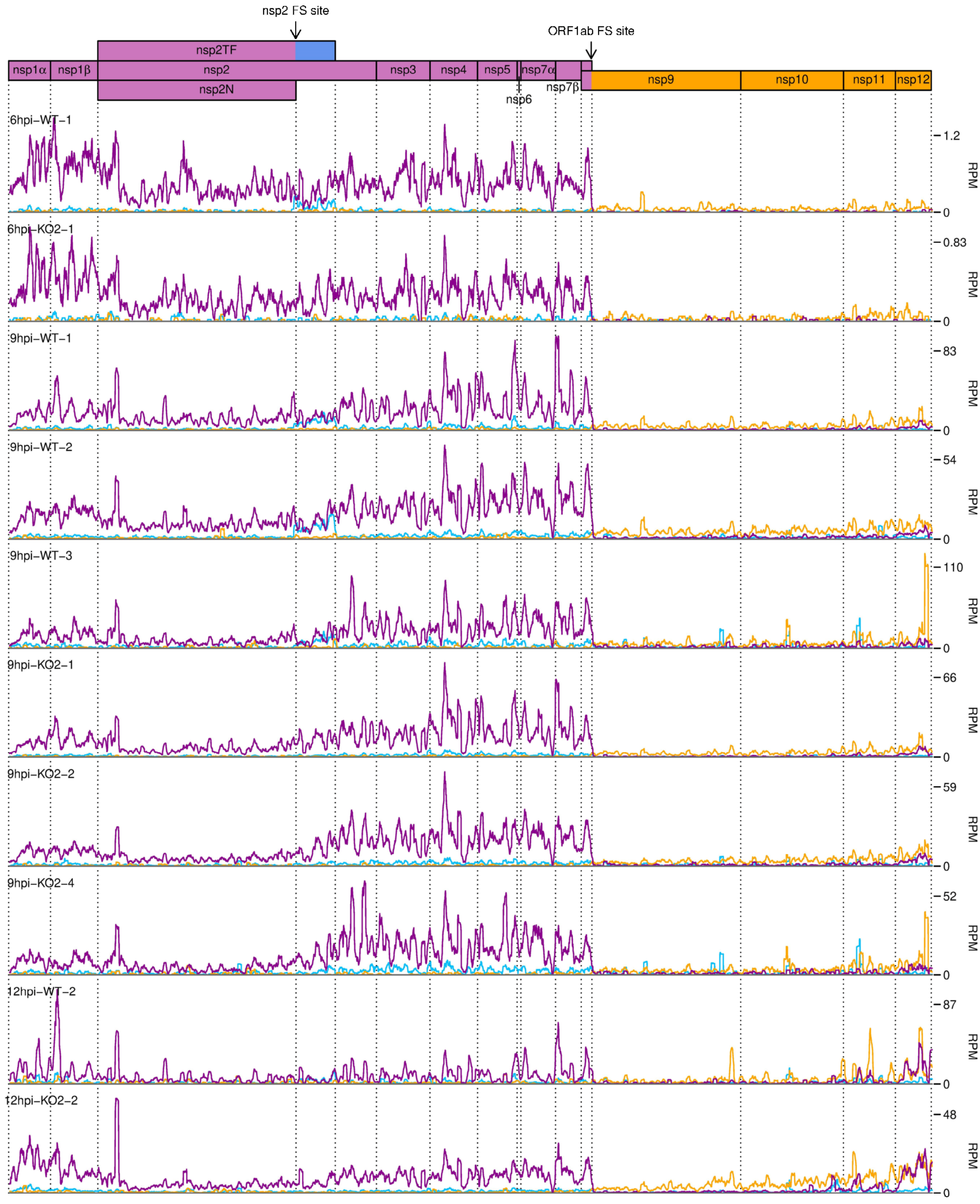


**Supplementary Figure 17. Distribution of RiboSeq reads in each phase in the ORF1ab region of the NA PRRSV genome, further replicates.**

Plot constructed as in Figure 10A.

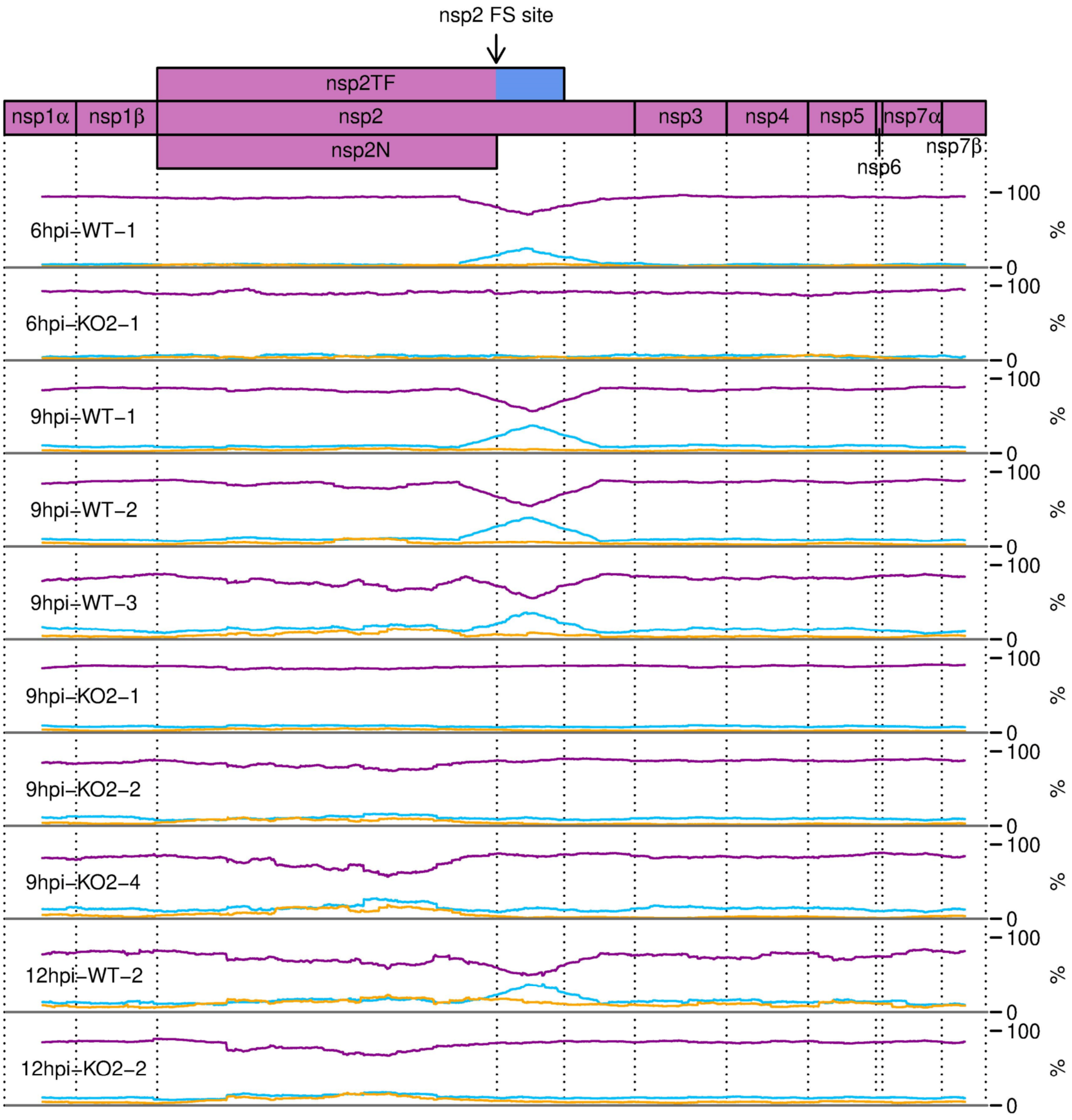


**Supplementary Figure 18. Percentage of RiboSeq reads in each phase across the ORF1a region of the NA PRRSV genome, further replicates.**

Plot constructed as in Figure 10B.

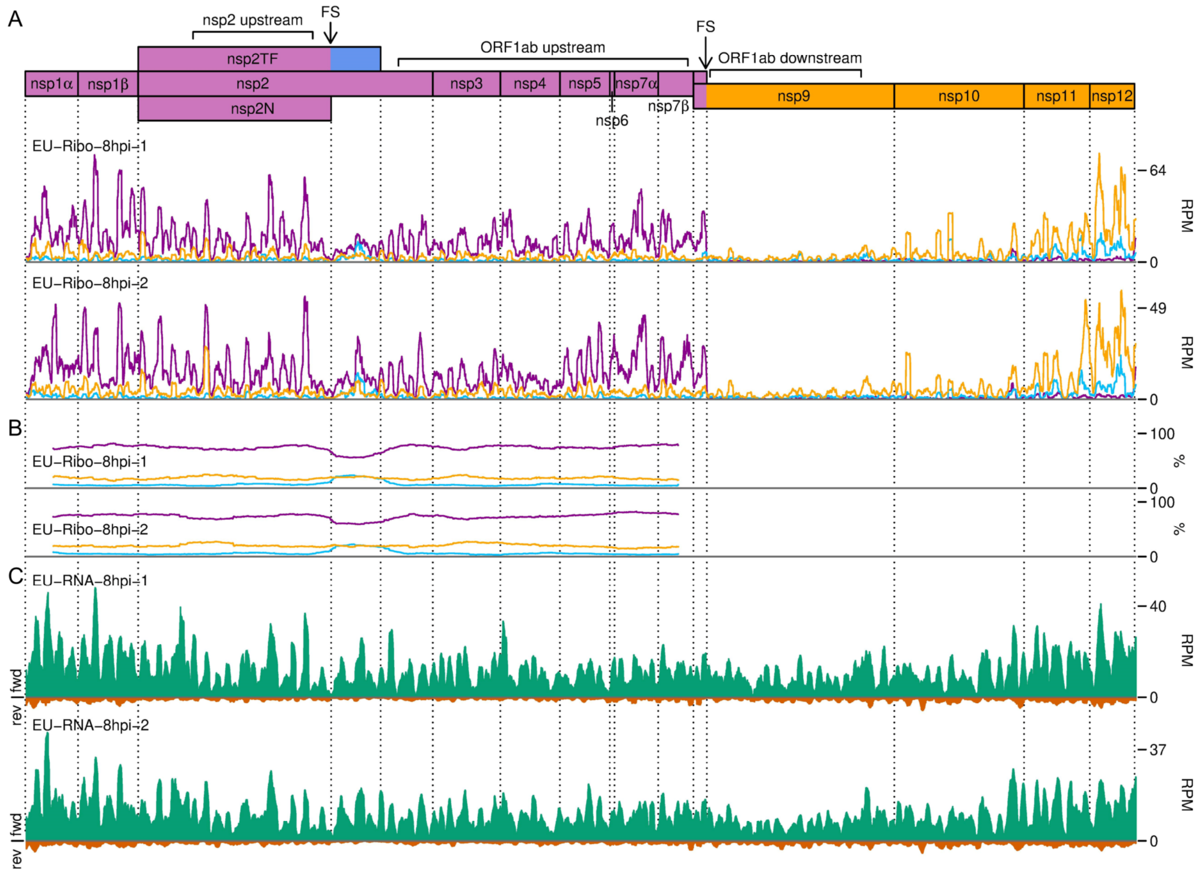


**Supplementary Figure 19. Distribution of reads in the ORF1ab region of the EU PRRSV genome.**

**A)** RiboSeq read densities separated according to phase. Plot constructed as described in Figure 10A, using read lengths with good phasing (indicated in Supplementary Figure 6D). **B)** Percentage of RiboSeq reads in each phase. Plot constructed as described in Figure 10B, using the same 183-codon running mean filter, and using read lengths with good phasing. **C)** RNASeq read densities. Plot constructed as in Figure 2B, with a 45-nt running mean filter applied, using all read lengths.

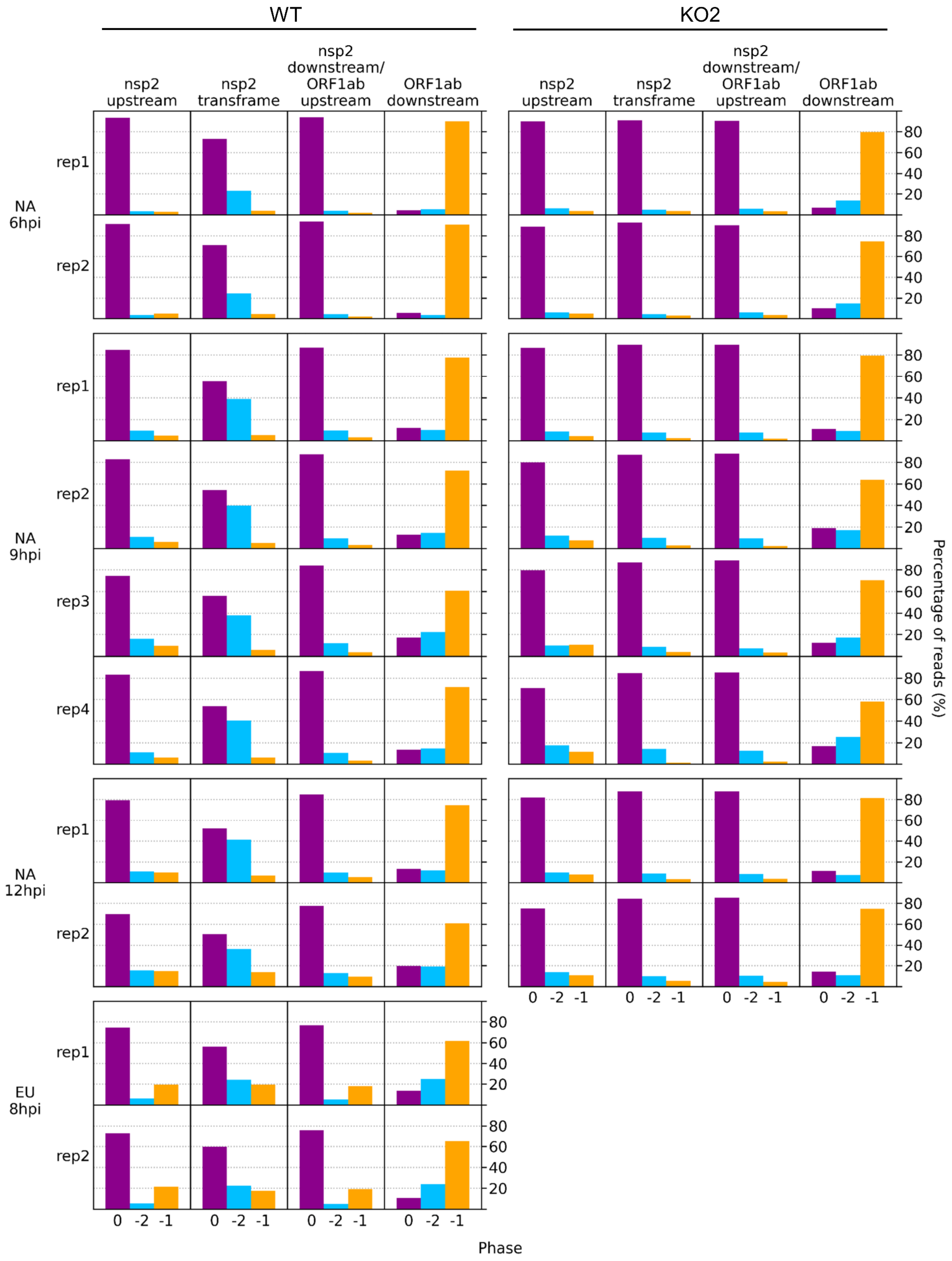


**Supplementary Figure 20. Bar charts of the percentage of reads in each phase in the specified regions of the viral genome.**

Read lengths showing good phasing were selected for inclusion in this analysis, as indicated in Supplementary Figure 4 for NA PRRSV libraries and Supplementary Figure 6D for EU PRRSV libraries. Genome regions are those annotated in Figure 10 and Supplementary Figure 19 (coordinates given in Supplementary Table 1).

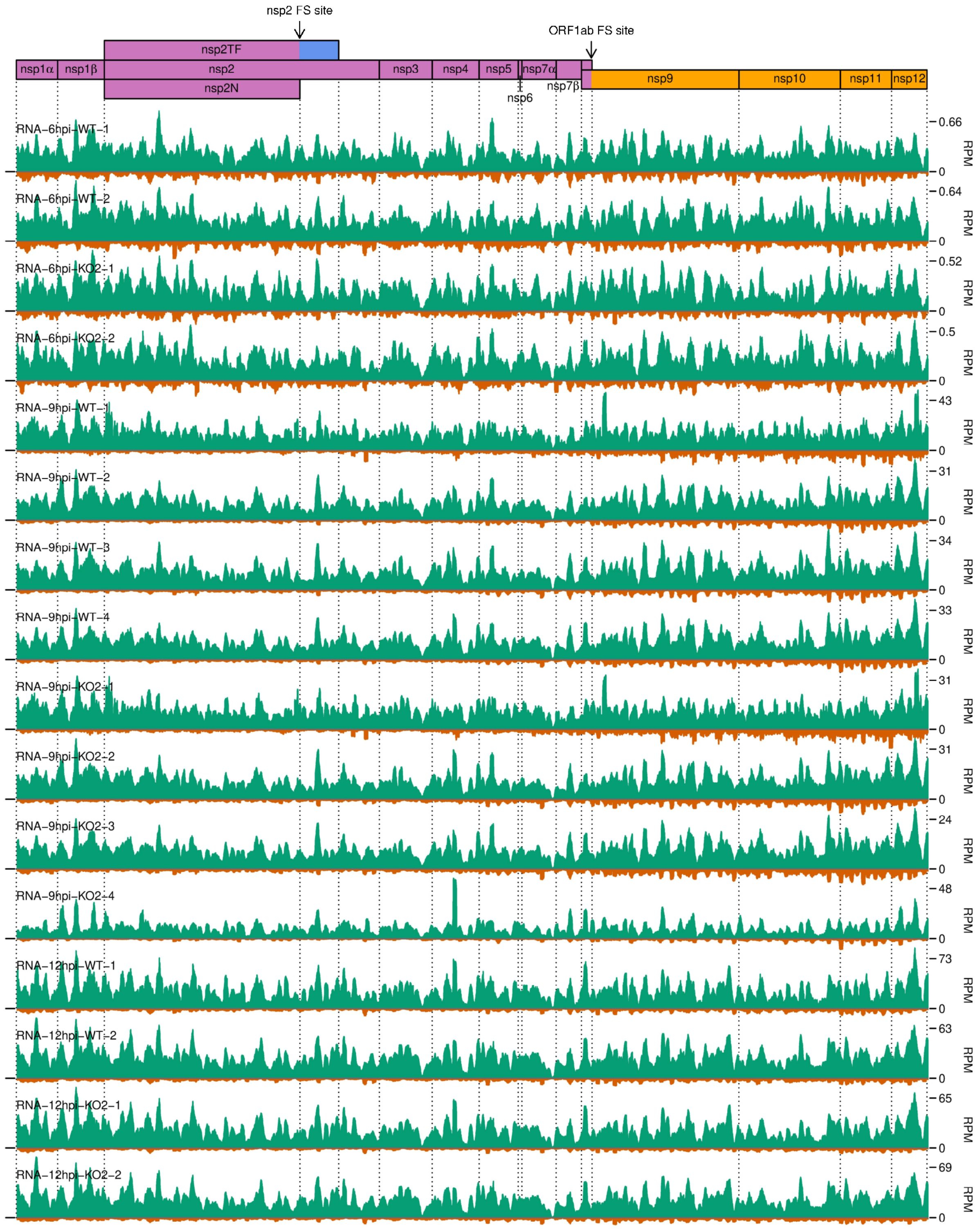


**Supplementary Figure 21. Distribution of RNASeq reads in the ORF1ab region of the NA PRRSV genome.**

Plot constructed as in Supplementary Figure 19C, using all read lengths.

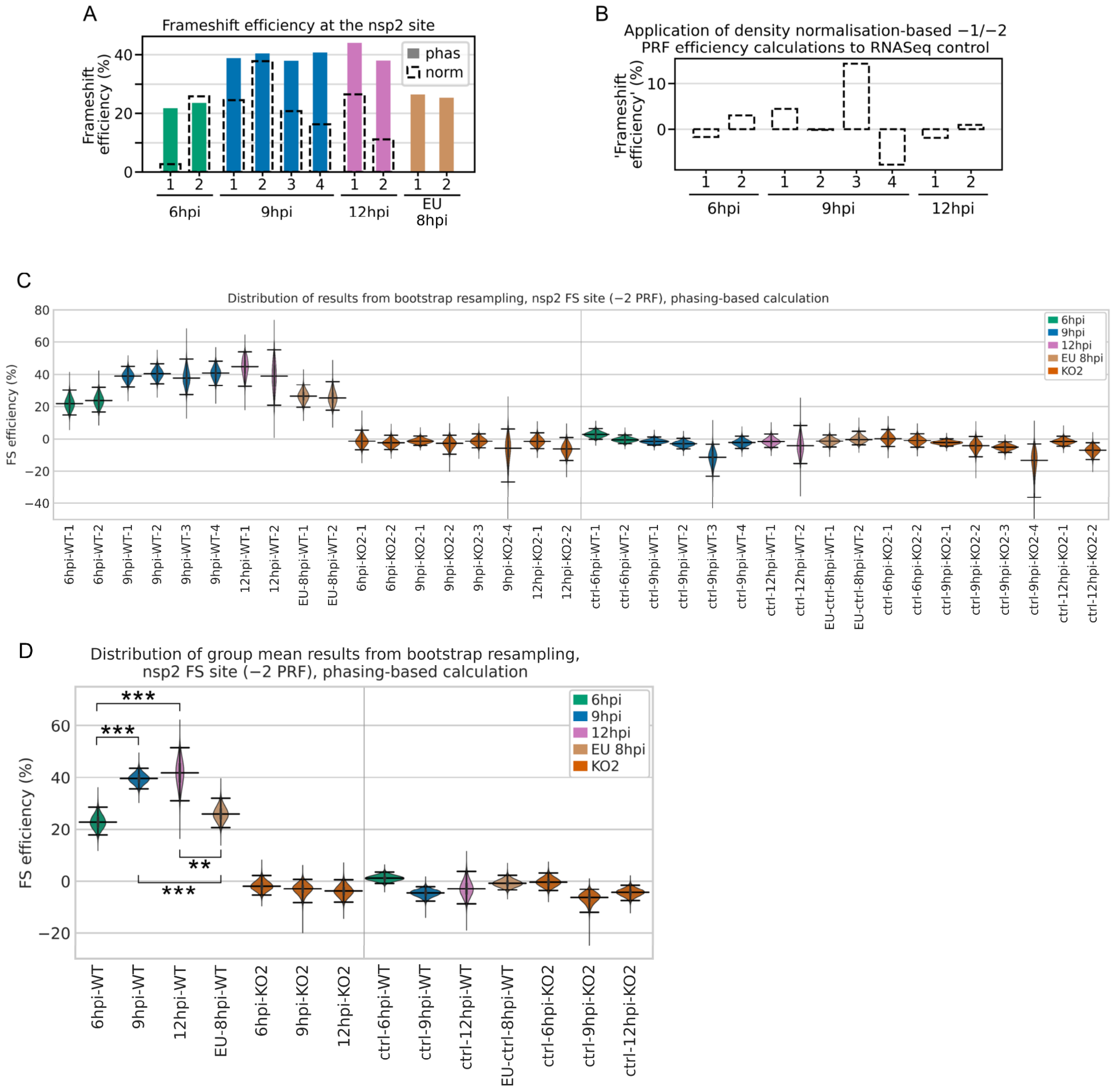


**Supplementary Figure 22. Bootstrapping and control analyses for nsp2 PRF efficiency.**

**A)** Comparison of the phasing-based (filled bars) and density normalisation-based (black dashed outlines) methods of PRF efficiency estimation at the nsp2 site. Note that the density normalisation-based method estimates the combined −1 and −2 PRF efficiency, whereas the phasing-based method estimates only −2 PRF efficiency. For EU PRRSV, no KO2 libraries were available so the normalisation-based method was not used. Only read lengths with minimal RNP contamination (NA PRRSV; Supplementary Figure 4) or good phasing (EU PRRSV; Supplementary Figure 6D) were used to perform these calculations. **B)** Application of the combined −1 and −2 PRF efficiency density normalisation-based calculation to the NA PRRSV RNASeq libraries as a negative control (expected value ~0%). All read lengths were used to perform these calculations. **C)** Distribution of results from calculation of −2 frameshift efficiency by the nsp2 phasing-based method after 100,000 bootstrap resamples of the codons in the upstream and transframe regions. The KO2 mutant libraries provide a negative control. As an additional negative control, the same calculations were performed using the downstream region instead of the transframe region, expected to yield ~0% frameshift efficiency (region coordinates given in Supplementary Table 1). To aid visibility, the y axis was truncated and the lower limits of the 9hpi-KO2-4 bootstrap populations are off scale. For all violin plots (here and in panel D), horizontal lines represent the median of the results distribution, and the 95% confidence intervals determined using the bias-corrected accelerated (BCa) method. **D)** Distribution of bootstrap results from calculation of mean −2 frameshift efficiency by the nsp2 phasing-based method. For each of the 100,000 resamples in panel C, the mean result was calculated for each indicated group of libraries, generating these distributions. Plot constructed as in panel C, using BCa 95% confidence intervals. Statistical significance of differences between non-control groups was determined using BCa confidence intervals (as described in Methods), with asterisks representing: * $p < 0.05$, ** $p < 0.005$; *** $p < 0.0005$.

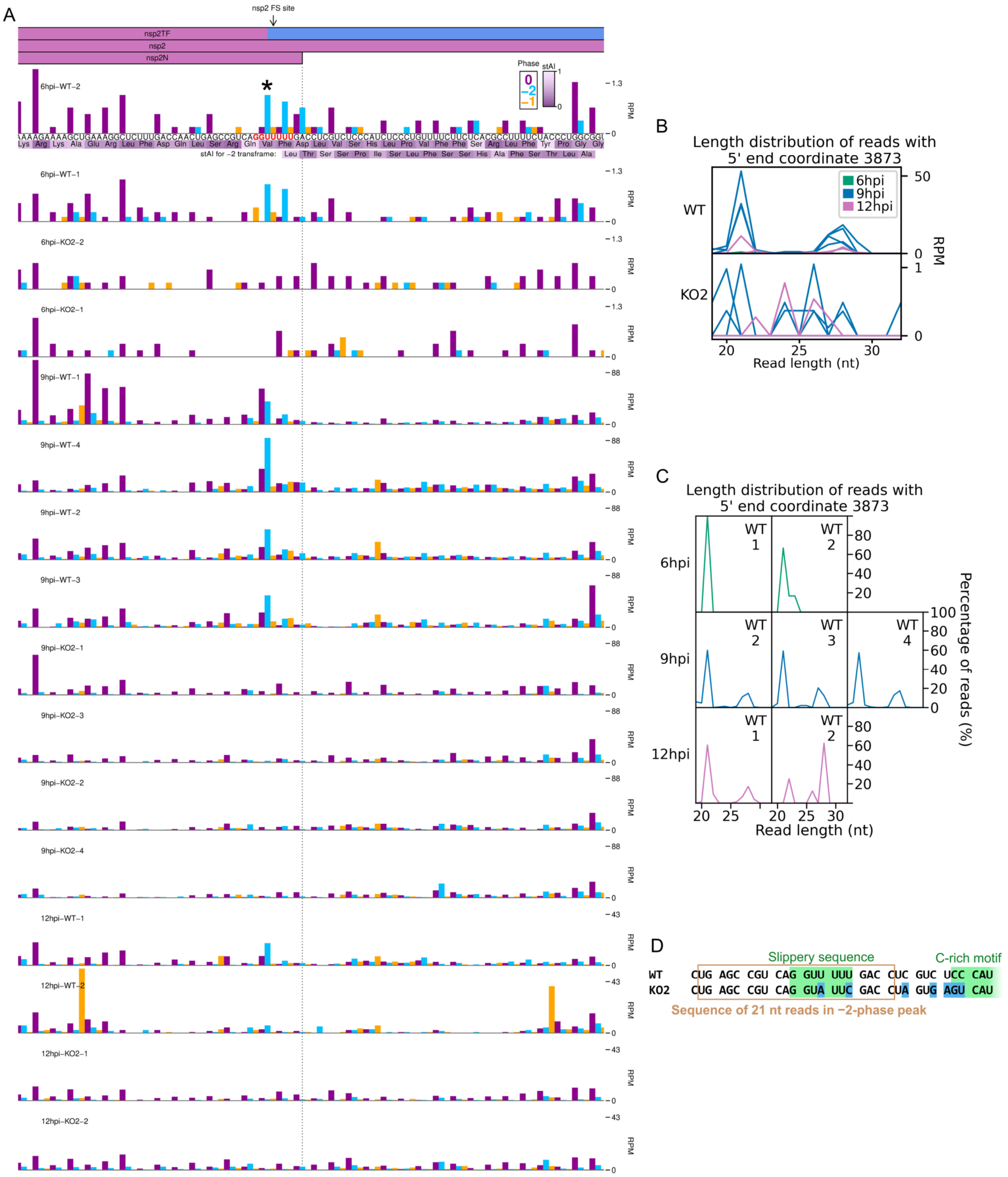


**Supplementary Figure 23. RiboSeq read densities around the nsp2 frameshift site.**

**A)** RiboSeq read densities around the nsp2 frameshift site. Plot constructed as in Figure 6D, with a heatmap of the species-specific tRNA adaptation index (stAI) of each codon underlaid beneath the amino acid identities (*Macaca mulatta* values were used as *C. sabaeus* values were unavailable). These values indicate the extent to which the codon is adapted to the host tRNA pool, with low values indicating poor adaptation, which may result in slow translation[1,2]. Codons in-frame with respect to nsp2 are displayed above those of the overlapping nsp2TF −2 frame. The slippery sequence is printed in red and the most prominent peak on the WT slippery sequence is indicated with an asterisk. As frameshifting could potentially affect the lengths of RPFs, all read lengths were used to make this plot. All libraries for a given timepoint are set to the same RPM scale on the y axis, with no running mean filter applied. **B)** Length distribution of reads that make up the blue peak marked with an asterisk in panel A. Length distribution is plotted as the number of reads (RPM) of each length, so libraries with a greater number of reads at this position (relative to total virus- and host-mapping positive-sense reads) appear more prominent. Due to considerable differences in read counts at this base between WT (top) and KO2 (bottom) these libraries were plotted on separate scales. Note that the very low number of reads at this position on the KO2 genome means the length distributions in the KO2 panel are highly subject to noise. **C)** Data for selected libraries from B reproduced to show the number of reads of each length as a percentage of the total number of reads mapping to this base (all libraries are therefore displayed on the same scale). KO2 libraries were omitted, and 9 hpi WT replicate 1 was omitted due to the exclusion of reads below 25 nt in the preparation of this library. **D)** Positions of mutations in the KO2 viral genome (blue) relative to the positions of reads comprising the majority of the peak marked with an asterisk in panel A (brown box).

### Supplementary Table legends

Supplementary Tables included as separate files.

**Supplementary Table 1. Coordinates of regions of the NA and EU PRRSV genomes used for each analysis.**

All coordinates denote the regions within which the 5′ end of the reads must map to qualify for inclusion in the analysis. **Sheet 1:** general plots and analyses (other than quantification of viral gene expression). **Sheet 2:** transcript abundance estimation. **Sheet 3:** translation level estimation.

**Supplementary Table 2. Composition of libraries.**

Number of reads assigned to each category. Reads were classified as "too short" if their inferred original fragment length was shorter than the minimum intended length experimentally purified (25 nt for all RNASeq libraries; for RiboSeq libraries: 25 nt for EU PRRSV libraries and NA PRRSV 9 hpi replicate one libraries, 19 nt for all other libraries).

**Supplementary Table 3. Junctions in the NA PRRSV transcriptome at 3 hpi.**

**Sheets 1 and 3 ("Filtered TRS junctions" and "Filtered non-TRS-spanning junctions")**: Final junctions after clustering and merging within each library and then filtering to select only junctions present in multiple replicates. All results are shown, including those with too few reads to pass the filter for inclusion on the sashimi plots. "Donor cluster" and "Acceptor cluster" columns show the group of genomic coordinates that formed the final merged cluster for the donor and acceptor of each junction. "Input junctions" gives the pairings (donor_acceptor) of these coordinates in the input junctions from the input libraries, and "Input junction read counts" gives the corresponding number of junction-spanning (JS) reads for each pairing. "Total junction read counts" is the sum of all the input junction read counts. "Donor midpoint" and "Acceptor midpoint" columns give the midpoints of the ranges of coordinates within the donor and acceptor clusters – these were used as the junction coordinates for making the sashimi plots and for calculating the number of non-junction-spanning reads spanning each site. The "Libraries with reads supporting junction" column gives the libraries in which the input junctions were found. The "Row IDs" column gives the row IDs of the input junctions from these libraries, which were merged to generate the output junction in this sheet. These row IDs can be used to inspect the input junctions from each individual library pre-merging, which can be found in the "Input TRS junctions" and "Input non-TRS-spanning junctions" sheets. The "sgRNA name" column (TRS junction sheet only) gives the name of the sgRNA, including non-canonical sgRNAs and minor transcript variants ("-" = not applicable). "Deletion length" (non-TRS-spanning sheet only) is the length, in nucleotides, of the region that is deleted due to each junction. "Donor non-JS read count" and "Acceptor non-JS read count" columns give the number of non-junction-spanning reads that span each site. "Proportion of JS reads at donor site" and "Proportion of JS reads at acceptor site" columns give the proportions of junction-spanning reads at each site, calculated as described in Methods. **Sheets 2 and 4 ("Input TRS junctions" and "Input non-TRS-spanning junctions")**: Junctions after clustering and merging within each individual library, which formed the input for the final filtering step that tests for presence in multiple replicates and merges matching junctions. Columns are as described for sheets 1 and 3.

**Supplementary Table 4. Junctions in the NA PRRSV transcriptome at 6 hpi.**

Table and columns as described for Supplementary Table 3.

**Supplementary Table 5. Junctions in the NA PRRSV transcriptome at 9 hpi.**

Table and columns as described for Supplementary Table 3.

**Supplementary Table 6. Junctions in the NA PRRSV transcriptome at 12 hpi.**

Table and columns as described for Supplementary Table 3.

**Supplementary Table 7. Junctions in the EU PRRSV transcriptome at 8 hpi.**

Table and columns as described for Supplementary Table 3.

**Supplementary Table 8. ORFs in the PRRSV translatome detected by PRICE.**

**Sheet 1:** Output from running PRICE on the NA PRRSV libraries, filtered to select viral ORFs with FDR-corrected $p$ value < 0.05. Columns and outputs are as described in the PRICE manual (output file ${prefix}.orfs.tsv), with two additional columns: "ORF name", giving the assigned ORF names, and "FDR-corrected p value", giving the $p$ value after Benjamini-Hochberg correction for multiple testing. Novel ORFs identified in this study are written in bold. Note that some canonical ORFs have differences in start codon designation compared to the reference genome annotation. **Sheet 2:** Output from running PRICE on the EU PRRSV libraries. Results were filtered to select viral ORFs with FDR-corrected *p* value < 0.05, and results for canonical ORFs with *p* values > 0.05 (E, 5a and nsp2TF transframe) were appended. Table as described for sheet 1.

**Supplementary Table 9. Host differential gene expression in WT vs mock at 12 hpi.**

**Sheets 1–3:** Differential transcription analysis. Significantly up-regulated and down-regulated genes are shown in sheets one (TS_up) and two (TS_down), respectively, while sheet three (TS_full) shows the results for all genes included in the analysis. The columns baseMean, log2FoldChange, lfcSE, stat, pvalue, and padj are as described in the DESeq2 documentation. The gene_ID column gives the ensembl gene ID, and the external_gene_name and wikigene_description columns give the name and description associated with these gene IDs. The p_adj_final column gives the final FDR-corrected *p* values that were used to make the volcano plots and to determine significance (while p_final gives the counterpart before correction for multiple testing). Where *p* value histograms were anti-conservative (WT vs mock and KO2 vs mock), p_adj_final and p_final are the same as padj and pvalue, respectively. For KO2 vs WT, the *p* value histogram was conservative, so *p* values were corrected using fdrtool to generate the values in p_adj_final and p_final (qval and pval from the fdrtool output, respectively). Columns entitled with library names contain the read counts for each gene in that library, generated using HTSeq and normalised for library size using DESeq2. **Sheets 4–6:** Differential TE analysis. Significantly up-regulated and down-regulated genes are shown in sheets four (TE_up) and five (TE_down), respectively, while sheet six (TE_full) shows the results for all genes included in the analysis. The columns log2FC_TE_final, pvalue_final, pvalue.adjust, log2FC_TE_v1, pvalue_v1, WT_log2TE, mock_log2TE, log2FC_TE_v2, and pvalue_v2 are as described in the xtail documentation (where log2FC_TE_final and pvalue.adjust are the final values used to make the volcano plots and determine significance). Columns entitled with library names contain the read counts for each gene in that library, generated using HTSeq and normalised for library size using DESeq2 (which formed the input for xtail). **Sheets 7–10:** GO term enrichment analysis. Lists of significantly differentially expressed genes from sheets one, two, four and five were used as input for DAVID to determine whether any GO terms associated with these genes were significantly enriched. All columns are as described in the DAVID documentation for "functional annotation chart report".

**Supplementary Table 10. Host differential gene expression in KO2 vs mock at 12 hpi.**

Table as described for Supplementary Table 9.

**Supplementary Table 11. Host differential gene expression in KO2 vs WT at 12 hpi.**

Table as described for Supplementary Table 9.
